# Development of difluoro-Kdn mechanism-based probes to label and visualize Kdnases in *Aspergillus fumigatus*

**DOI:** 10.64898/2026.03.12.711403

**Authors:** Erianna I. Alvarado-Melendez, Jacq van Neer, Hans de Cock, Tom Wennekes

## Abstract

Kdnases have been reported in a variety of organisms, including marine species such as trout and oysters, the opportunistic Gram-negative bacterium *Sphingobacterium multivorum*, and several fungal species of the genus *Aspergillus*, including *Aspergillus terreus* and *Aspergillus fumigatus*.. In particular, the Kdnase from the opportunistic airborne pathogen *Aspergillus fumigatus* (*Af*Kdnase) plays an important role in fungal cell wall integrity and virulence, although the underlying mechanisms remain unclear. To better understand this class of enzymes, selective and sensitive tools are required for discovery, detection and visualization of active Kdnases in complex biological samples. In this work, we report the development of difluoro-Kdn mechanism-based probes functionalized with azide and biotin tags for labeling and detection of Kdnases. We show that the probes exhibit selectivity for Kdnase over the neuraminidases tested and efficiently label recombinantly expressed *Af*Kdnase at micromolar concentrations. In addition, using the azide-bearing probe and click chemistry, we successfully visualized native Kdnases in *A. fumigatus* mycelia, demonstrating their utility for studying these enzymes in crude biological samples and highlighting their potential for discovering Kdnases in other organisms including fungal and bacterial species.

## Introduction

The monosaccharide, 2-keto-3-deoxy-D-glycero-D-galacto-nononic acid, known as Kdn, is a naturally occurring sialic acid. It is an analogue of the key human sialic acid, *N*-acetylneuraminic acid (Neu5Ac), differing by the absence of an *N*-acetamido group at C5. Kdn occurs in glycoconjugates from diverse sources, including marine vertebrates, fungi, bacteria, and mammals. In mammalian cells, Kdn occurs both as free monosaccharide and poly-Kdn, and as a replacement of Neu5Ac on the terminal position in glycolipids and glycoproteins, though at levels 100 to 1000 times lower than Neu5Ac. (1) In humans, Kdn is found in lung epithelium, ovarian cancer, and red blood cells, predominantly as α2,8-linked poly-Kdn. (2)

Terminal Kdn can be hydrolyzed from glycoconjugates by specialized exo-sialidases known as Kdnases. These enzymes have been identified in fish species, (3, 4) a Gram-negative opportunistic bacterium *Sphingobacterium multivorum* (5, 6) and marine invertebrates (7) In 2010, the discovery of a retaining Kdnase from *Aspergillus fumigatus* (*Af*Kdnase) (2, 8, 9) extended Kdnase biology to the fungal kingdom. *A. fumigatus* is an opportunistic airborne pathogen and the leading cause of invasive aspergillosis (IA), a frequently fatal infection in immunocompromised patients. Treatment options are constrained by drug toxicity (10) and rising of fungal resistance. (11) Since *Af*Kdnase is essential for fungal cell wall integrity, nutrient acquisition, and pathogenicity, (9) and humans lack Kdnase homologues, this represents a potential attractive and selective target for antifungal therapeutic interventions.

However, despite their relevance, Kdnases remain underexplored. Tools to detect, visualize, and profile these enzymes are scarce. Previous studies by Bennet and co-workers characterized fungal Kdnases, using the inhibitor 2,3-diF-Kdn, and indirectly mapped their localization at the cell surface with a fluorescent benzothiazolyl-4-bromophenyl Kdn probe (see **Scheme 1A**). (12) While informative, this approach provides only indirect localization evidence. Beyond structural and localization studies, Bennet’s group has developed strategies to screen for Kdnase activity using Kdn thioglycosides (13) and a self-immolative Kdn glycoside for high-throughput screening of Kdnase inhibitors. Using this strategy they found two non-sugar competitive inhibitors with IC_50_ values of 23 and 118 μM. (14) Around the same time, Scalabrini et al. reported a multivalent, non-covalent *Af*Kdnase inhibitor with an IC_50_ of 1.5 μM, the most potent inhibitor reported to date (see **Scheme 1A**). (15)

To expand the chemical toolbox for Kdnase inhibition and enable direct and selective detection of retaining Kdnases in biological samples, in this work we developed difluoro-Kdn mechanism-based probes. This class of probes was first introduced by Withers et al. and has since been widely used for screening of retaining glucosidases, neuraminidases and fucosidases. (16–19) Retaining glycosidases operate via a two-step active-site mechanism. Following substrate binding, a covalent glycosyl-enzyme intermediate is formed (Scheme 1B), resulting in cleavage of the glycosidic bond and release of the aglycone. In the second step, hydrolysis of this intermediate regenerates the active enzyme. Our design features a Kdn-based scaffold bearing a C9 azide for click-chemistry detection or affinity tagging, together with two strategically positioned fluorine atoms (**Scheme 1C**). Fluorination at C3 destabilizes the oxocarbenium-like transition state involved in both catalytic steps. Fluorination at C2 provides a suitably reactive leaving group that permits formation of the covalent intermediate, while significantly slowing or preventing the subsequent hydrolysis step. This traps the covalent probe–enzyme intermediate, thereby enabling irreversible enzyme labeling and inactivation.

In this work, we show that our probes covalently label Kdnases, enabling both inhibition and specific labeling of *Af*Kdnases. We provide the first direct visualization of these enzymes in *A. fumigatus* mycelia (see Scheme 1D), demonstrating the utility of the probes for their use in complex native samples. We envision that these chemical tools can be used for the discovery and characterization of other retaining Kdnases across fungi and other organisms, including bacterial species. Ultimately, this approach could advance our understanding of Kdnase biology and shed light on the role of fungal Kdnases in pathogenesis.

**Scheme 1.**
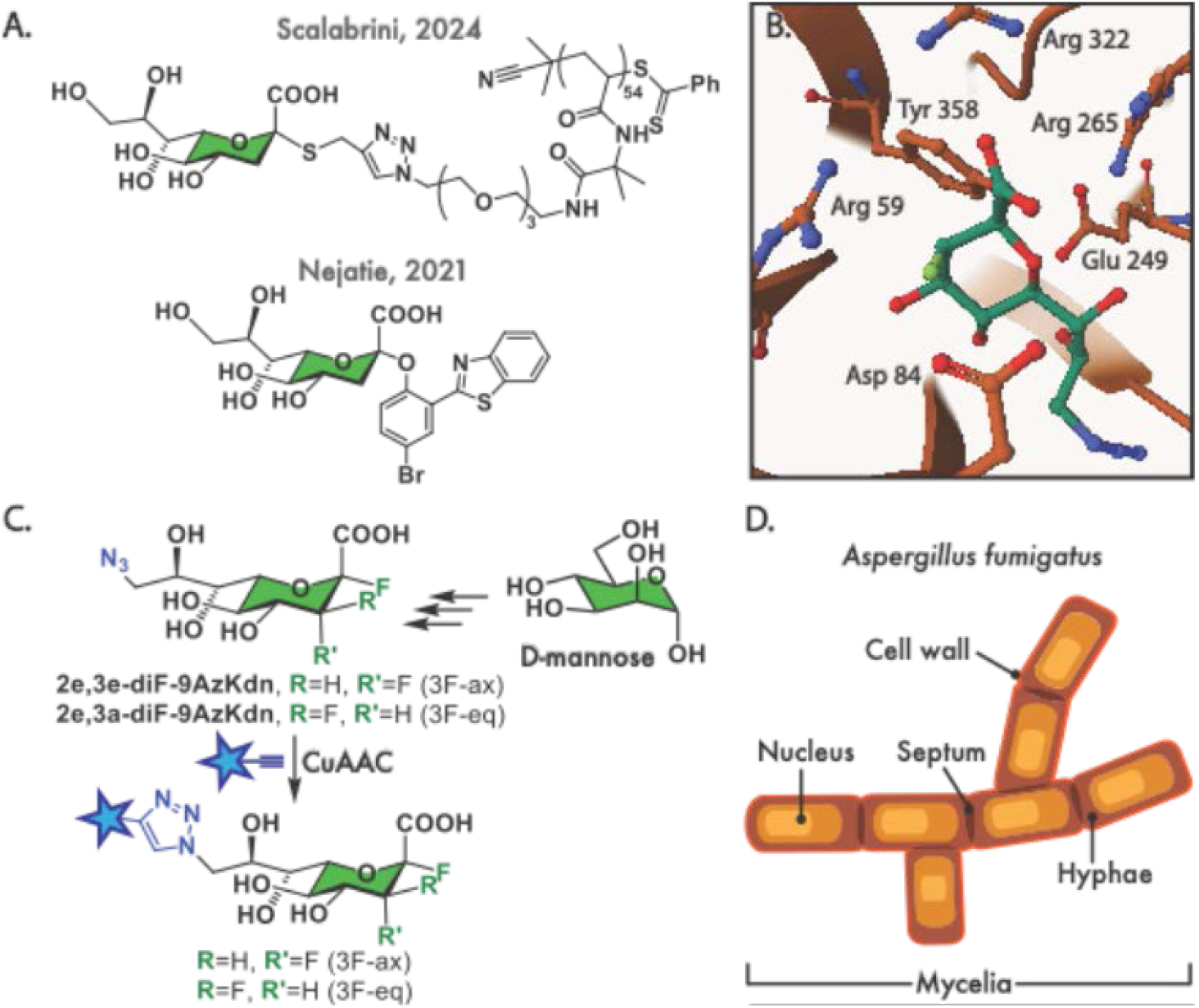
**(A)** Structure of the most potent reversible inhibitor of *Af*Kdnase (Scalabrini, 2024), and the imaging fluorescence probe (Nejatie, 2021) described in previously. **(B)** Enzyme-probe intermediate between the *Aspergillus fumigatus* Kdnase (*Af*Kdnase) and **2e**,**3a-diF-9AzKdn. 2e**,**3a-diF-9AzKdn** was docked into the active site of *Af*Kdnase structure (PDB: 2X2J) using Yasara and Mol* viewer for 3D visualization. The catalytic residues are highlighted and labeled in the structure, where Tyr 358 is the nucleophile and forms a covalent bond with the probe. **(C)** Probes designed and developed in this work. **(D)** Structure of *Aspergillus fumigatus* Mycelia

## Results

### Synthesis of the difluoro-Kdn probes

We synthesized four mechanism-based Kdn probes that vary in reactivity and functional handle (Figure 1). Two probes incorporate an azide group for click-chemistry applications (**2e**,**3a-diF-9AzKdn** and **2e**,**3e-diF-9AzKdn**), while two are functionalized with biotin (**2e**,**3a-diF-9bKdn** and **2e**,**3e-diF-9bKdn**). To investigate the effect of fluorine configuration at the C3 position on Kdnase inactivation, we designed these probes with either an axial (3a) or equatorial (3e) fluorine atom at C3. This design was informed by previous studies demonstrating that axial and equatorial C3 difluoro substitutions differentially modulate the inactivation and reactivation kinetics of sialidase probes. (20, 21)

**Figure 1.**
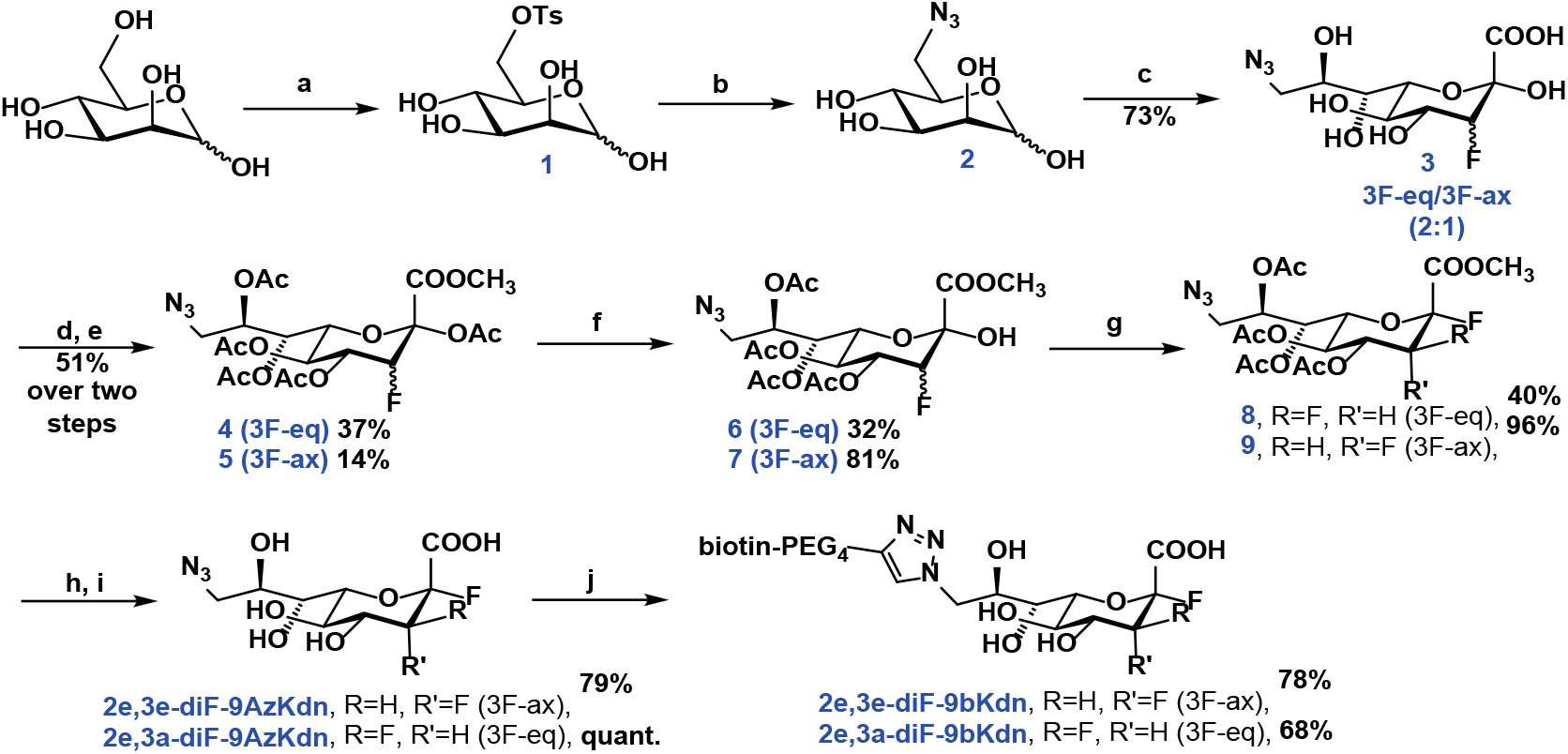
Synthesis of Kdn ABPs. Reagents and conditions: (a) TsCl, *py*, RT, 18 h; (b) NaN_3_ (acetone:H_2_O, 3:1), 50 °C; (c) NANA aldolase, βF-pyruvate, H_2_O, RT, 6 days; (d) TFA, CH_3_OH, RT, o/n; (e) Ac_2_O, pyridine, RT, o/n; (f) hydrazine acetate, CH_2_Cl_2_/CH_3_OH, 0 °C, 7 h; (g) DAST, CH_2_Cl_2_, −30 °C, 1-2 h; (h) Na_2_CO_3_, CH_3_OH, RT, 1 h; (i) NaOH (aq.), pH 11, RT, 1 h; (j) alkyne-PEG_4_-biotin, Cu-THPTA, Na-L-ascorbate, H_2_O, RT, 18h.

**Figure 2.** Inhibition of *Af*Kdnase. We determined IC_50_ values using the fluorometric assay with 4MU-Kdn. For this, we incubated *Af*Kdnase with varying concentrations of the probes and calculated IC_50_ values using nonlinear regression analysis in GraphPad Prism.

We first synthesized 6-azido-D-mannose via a previously reported procedure. (22) We used the *N*-Acetylneuraminic acid aldolase (Neu5Ac aldolase, EC: 4.1.3.3) to catalyze the synthesis of 3F-Kdn from 6-azido-D-mannose and 3-fluoropyruvate. 3F-Kdn (a mixture of axial and equatorial 1:2) was converted to the methyl ester using TFA in methanol and acetylated with acetic anhydride in pyridine to give compounds **4** and **5**. Selective deacetylation at the anomeric position was achieved using hydrazine acetate, yielding the hemi-acetals **4** and **5**. Compounds **4** and **5** were treated with diethylaminosulfur trifluoride (DAST) at −30 °C to give the α-fluorides **8** and **9**. Subsequent deprotection yielded probes **2e**,**3a-diF-9AzKdn** and **2e**,**3e-diF-9AzKdn**. Finally, copper-catalyzed azide-alkyne cycloaddition (CuAAC) reaction was used to conjugate alkyne-PEG_4_-biotin, affording probes **2e**,**3a-diF-9bKdn** and **2e**,**3e-diF-9bKdn**. All intermediates were characterized by ^1^H-NMR, ^13^C-NMR, ^19^F-NMR and ESI-MS. The synthesis and the characterization data can be found in supporting information S1.

### Enzymatic activity and inhibition of recombinantly expressed *Af*Kdnase

We investigated the ability of the probes to inhibit the recombinantly expressed *Af*Kdnase from *Aspergillus fumigatus* Af293. To monitor the Kdnase activity, we used the fluorogenic substrate 4-methylumbelliferyl-Kdn (4MU-Kdn) and measured the fluorescence of the released 4-methylumbelliferone. We verified that *Af*Kdnase did not exhibit sialidase activity by using MUNANA (4-methylumbelliferyl-*N*-acetyl-α-D-neuraminic acid) as the fluorescent substrate. We determined the Michaelis-Menten parameter K_*m*_ of the *Af*Kdnase using 4MU-Kdn, and found a value of 0.27 mM, which is consistent with the K_*m*_ previously reported for this enzyme (0.23 ± 0.02 mM). (2) (see supporting information S2).

We then evaluated the inhibitory ability of the probes. For this assay, we incubated the enzyme with different concentrations of the probes and followed the enzymatic activity using 4MU-Kdn. We plotted fluorescence intensity against probe concentration to calculate the apparent IC_50_ values for each probe using nonlinear regression analysis in GraphPad Prism (see **Figure 1**). Probes bearing the fluorine atom in the axial configuration showed an increased inhibitory activity against *Af*Kdnase compared to their 3F(equatorial)-counterparts. Click of alkyne-PEG_4_-biotin to the C9-N_3_ led to a decrease in inhibitory activity. Among the probes tested in this work, the **2e**,**3a-diF-9AzKdn** was the most selective and sensitive inhibitor of the *Af*Kdnase.

### Time-course study for *Af*Kdnase reactivation after incubation with probe 2e,3a-diF-9AzKdn

One limitation of difluoro mechanism-based probes is the eventual hydrolysis of the covalent intermediate formed with the target glycosyl hydrolase. To assess the stability of this covalent intermediate, and therefore the feasibility of using **2e**,**3a-diF-9AzKdn** for labeling *Af*Kdnases, we performed a time-course experiment to monitor enzyme reactivation. In this assay, recombinant His-tagged *Af*Kdnases were immobilized on Ni-NTA beads, the immobilized enzymes were inactivated with probe **2e**,**3a-diF-9AzKdn**, and washed to remove unbound probe. Aliquots of the inactivated *Af*Kdnase-beads suspension were then tested at defined time points for residual catalytic activity using 4MU-Kdn as the substrate. We observed progressive recovery of hydrolytic activity over time, reaching full reactivation after approximately seven hours (**Figure 3**).

**Figure 3.**
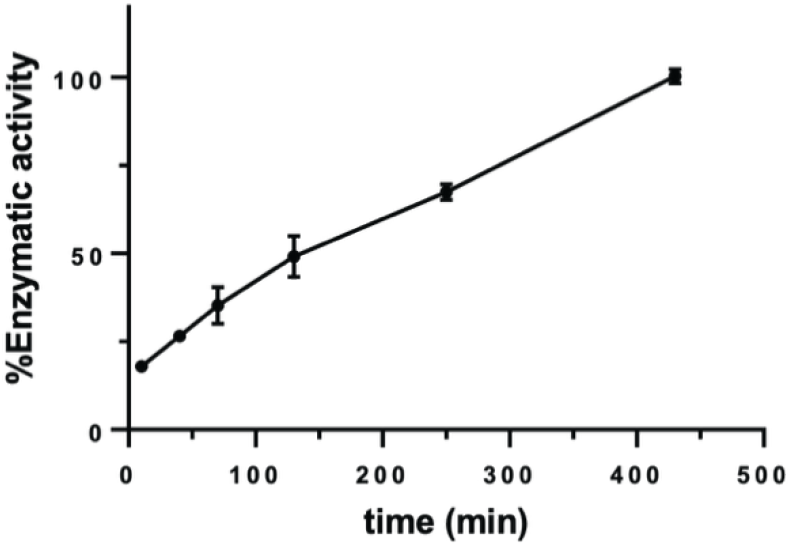
Reactivation of *Af*Kdnase after incubation with **2e**,**3a-diF-9AzKdn**. Kdnase activity was assayed at each time point using 4MU-Kdn.

### Labeling and detection of the recombinant *Af*Kdnase

To investigate the ability of the biotinylated probes (**2e**,**3a-diF-9bKdn** and **2e**,**3e-diF-9bKdn**) to covalently label *Af*Kdnase, we incubated the recombinantly expressed enzyme with each probe for one hour at room temperature and detected labeled proteins by Western blot using streptavidin-HRP (see **Figure 4**). Under these conditions, we found that **2e**,**3a-diF-9bKdn** was more sensitive in labeling *Af*Kdnase, with a visible band when used at 1.0 µM concentration. Probe **2e**,**3e-diF-9bKdn** only shows visible labeling when used at a concentration of 100 µM. These results are consistent with the inhibition assays. To test the selectivity of the probes, we used two bacterial neuraminidases *Pt*NanH1 and *Pt*NanH2. The results demonstrate that **2e**,**3a-diF-9bKdn** is selective towards *Af*Kdnase. We also used a neuraminidase ABP (**2**,**3-diF-Neu9biotin**) to show that *Af*Kdnase does not metabolize neuraminic acid analogs, as we saw in the activity assays.

**Figure 4.**
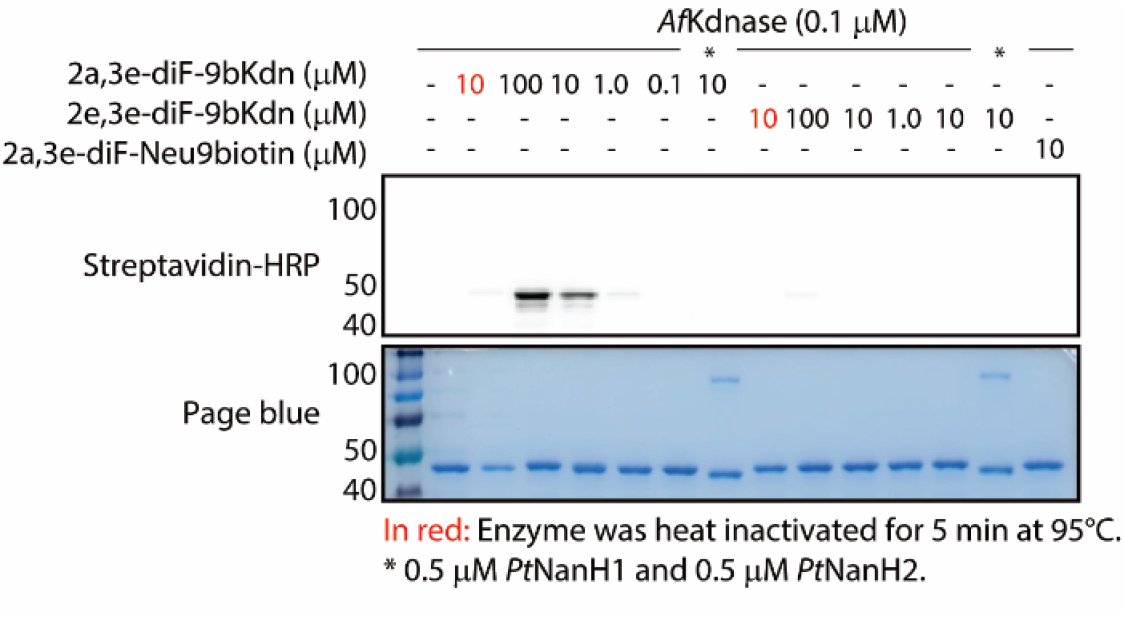
Labeling of recombinant *Af*Kdnase (44 kDa) with **2e**,**3a-diF-9bKdn** and **2e**,**3e-diF-9bKdn** at different concentrations. Recombinant neuraminidases *Pt*NanH1 (42 kDa) and *Pt*NanH2 (110 kDa) were used to test the selectivity of the probes towards Kdnases. The recombinant *Af*Kdnase is not labeled by the neuraminidase mechanism-based probe **2e**,**3a-diF-Neu9biotin**. Full size gels can be found in the supporting information S3.

### Inhibition of Kdnase activity in *A. fumigatus* mycelia

We next evaluated whether probe **2e**,**3a-diF-9AzKdn**, the best inhibitor against *Af*Kdnase, could inhibit Kdnase activity in *A. fumigatus in vitro*. First, we assessed Kdnase activity in both spores and mycelia of four *A. fumigatus* strains (Af293, CEA10, ATCC46645 and A1258) grown in minimal medium and in supplemented medium (0.3%, w/v, yeast extract and malt extract; 0.5%, w/v, peptone; and 1%, w/v, glucose). We detected Kdnase activity only in mycelia cultured for three days in the supplemented medium. We then tested the inhibitory effect of probe **2e**,**3a-diF-9AzKdn**. To this end, we pre-incubated the active mycelia with the probe for one hour at room temperature, followed by an *in vitro* Kdnase activity assay using 4MU-Kdn. We registered a marked decrease in Kdnase activity in the samples treated with **2e**,**3a-diF-9AzKdn** as shown in the graph below (**Figure 5**).

**Figure 5.** Fluorometric assay monitoring Kdnase activity in different *A. fumigatus* strains. We incubated the mycelia with the fluorogenic substrate 4-methylumbelliferyl-Kdn (4MU-Kdn) and measured the fluorescence of the released 4-methylumbelliferone after 150 min incubation at RT. The graph shows results from two biological replicates. Controls can be found in the supporting information S4.

### Visualization of Kdnase in *A. fumigatus* mycelia by fluorescence microscopy

Having confirmed that *A. fumigatus* mycelia exhibit Kdnase activity and having an indication that **2e**,**3a-diF-9AzKdn** can inhibit this activity, we next aimed to use the probe to visualize Kdnase enzymes in the mycelia. For this, we grew the *A. fumigatus* ATCC46645 for three days in supplemented medium, and then we incubated the mycelium with **2e**,**3a-diF-9AzKdn** for 30 minutes at RT. Following incubation, we performed a CuAAC reaction to click the AF_488_ fluorophore, and we visualized the labeled mycelia by fluorescence microscopy. To show that the labeling observed was due to the probe covalently attached to the enzyme, we used a control of mycelia incubated with only the click reaction mixture, along with a negative control (**Figure 6**). In the sample treated with **2e**,**3a-diF-9AzKdn** and the click mixture, we observed uniform or patchy green labeling of the hyphae with intensities which varies between hyphae detected in the microscopic images (**Figure 6A-C**), but appears to label the surface of the hyphae (**Figure 6B**); this fluorescence was not detected in the controls. Septa detected in the images appear not labeled by the probe (**Figure 6B** red arrows).

**Figure 6.**
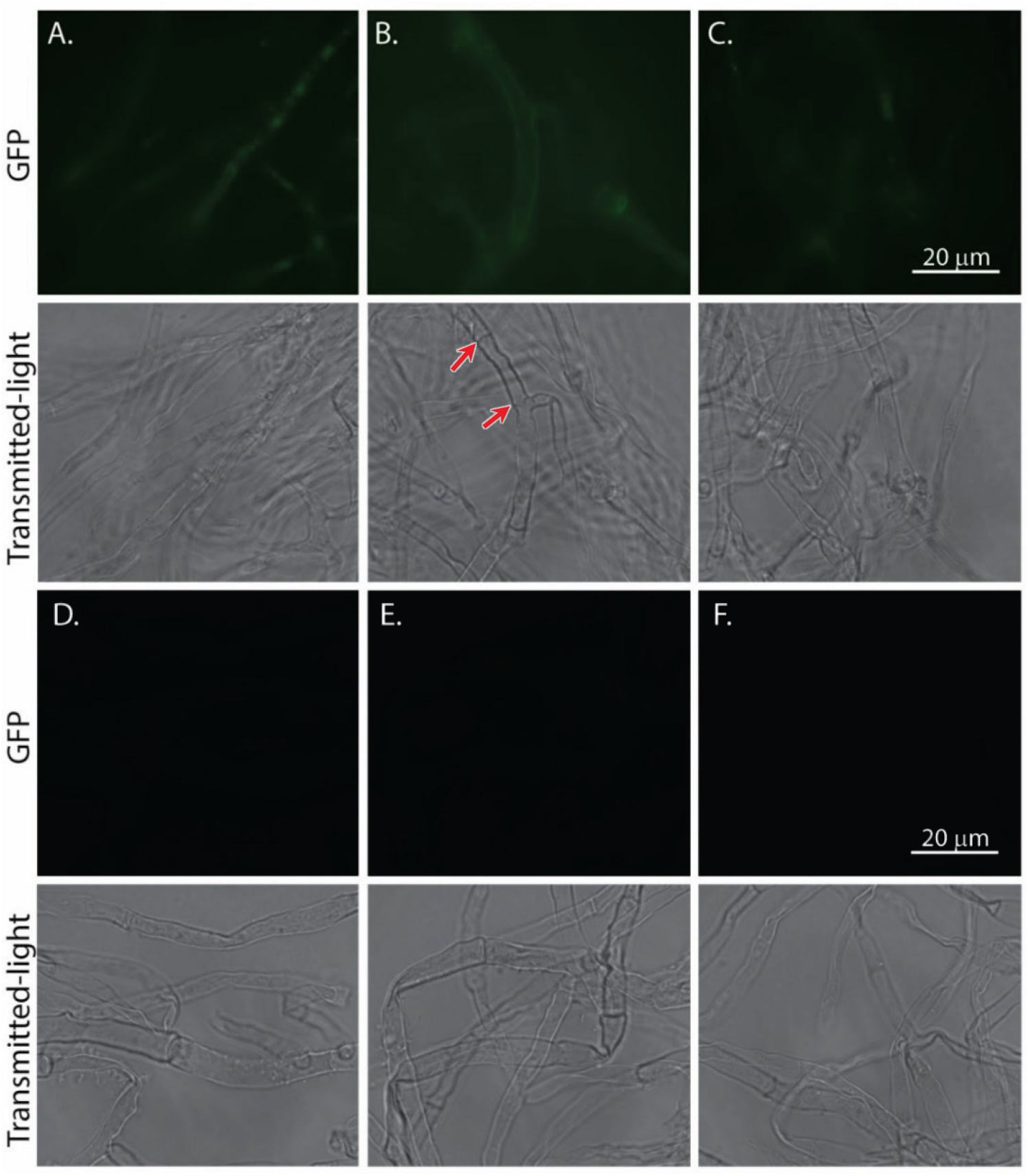
Microscopy image of *A. fumigatus* ATCC46645 hyphae. **(A-C)** Hyphae fluorescently labeled with **2e**,**3a-diF-9AzKdn** followed by CuAAC reaction with alkyne-AF_488_ (GFP and transmitted-light channels). **(D-E)** Hyphae treated and incubated with alkyne-AF_488_ (non-specific labeling control). **(F)** Unlabeled hyphae.

## Discussion

Kdnases have been detected in aquatic species such as trout and oyster, opportunistic fungi and bacteria, but not in humans. In humans, Kdn is present at only 0.1-1% of Neu5Ac levels. Interestingly, higher Kdn-to-Neu5Ac ratios have been observed in tumor tissues, including ovarian and lung cancer cells. (23, 24) This makes Kdnases an intriguing class of enzymes to explore. Additionally, the widespread occurrence of Kdn in mammals, fungi, and bacteria suggests that more, yet-undiscovered Kdnases may exist. Detecting and characterizing these enzymes is a crucial step toward understanding their roles in organismal survival and virulence.

Kdnase activity can be measured using a fluorometric assay with 4-methylumbelliferyl 3-deoxy-d-glycero-d-galacto-non-2-ulopyranosidonic acid (4MU-Kdn) as the substrate. However, this fluorescent substrate has previously been shown to be hydrolytically labile. (13) We observed that hydrolysis occurs even at neutral pH and is further accelerated at pH 4, the optimal pH for *Af*Kdnase, leading to high background fluorescence. These limitations make the assay unsuitable for long-term experiments or for sensitive detection of activity. Previous attempts to overcome this limitation using Kdn thioglycosides resulted in substrates that were less efficiently hydrolyzed by *Af*Kdnase. (13) Subsequently, self-immolative Kdn substrates were developed for high-throughput screening of Kdnase inhibitors. (14) This strategy is effective for screening Kdnase activity, but it only provides information on the catalytic activity of the enzyme.

In this work we describe the development of mechanism-based probes that can be used for direct and robust detection, visualization, and inhibition of Kdnases in biological samples. We based our design on the known covalent inhibitor **2e**,**3a-diF-Kdn** and introduced azide and biotin tags at the C9 position for detection. We synthesized difluoro-Kdn probes with the fluorine atom at C3 in both axial and equatorial configurations to evaluate how stereochemistry affects labeling and inhibitory activity.

Our inhibition assays demonstrate that probes bearing the fluorine atom in the axial configuration exhibit higher inhibitory activity against *Af*Kdnase compared to their equatorial C3 counterparts. Notably, the click modification of the C9 azide with alkyne-PEG_4_-biotin led to a reduction in inhibitory potency, likely due to steric effects or altered interactions within the active site. Among the probes tested, **2e**,**3a-diF-9AzKdn** is the most selective and sensitive inhibitor of *Af*Kdnase. A key limitation of difluoro-based mechanism probes is that the covalent intermediate they form with the enzyme hydrolyzes over time. To account for this, we designed experiments to monitor enzyme activity and labeling over extended periods. Importantly, we observed that the enzyme undergoes full reactivation within seven hours. This observation was considered when designing experiments for long-term inhibition and labeling.

In our labeling experiments, we found that probe **2e**,**3a-diF-9bKdn** detected *Af*Kdnase more sensitively than **2e**,**3e-diF-9bKdn**, consistent with the IC_50_ values observed. Similar trends have been reported for difluoro-Neu5Ac probes used in *Clostridium perfringens* and *Vibrio cholerae*. (19) This study showed that neuraminic acid probes with an equatorial C3 fluorine are less potent inhibitors of neuraminidases. In contrast, axial C3 fluorine forms favorable interactions via hydrogen bonds with enzyme residues, which likely explains the superior performance of the axial C3 probe in both labeling and inhibition.

We next moved to a more complex system and sought to fluorescently label Kdnases in mycelia samples to study the enzyme’s localization. Since Kdnase activity was detected in vitro only in mycelium grown in rich medium, mycelia cultivated under these conditions were used for the fluorescent labeling experiments. Using probe **2e**,**3a-diF-9AzKdn** followed by CuAAC reaction with alkyne-AF_488_, we detected Kdnases in live *A. fumigatus* cells, with no fluorescence observed in the control samples (see **Figure 6**). The fluorescence localizes primarily to the hyphae and appears to be localized at the surface, the expected location of this extracellular cell wall located enzyme. (12) Previously, Nejatie et al. (2021) used an indirect approach to localize Kdnases in *A. fumigatus* mycelia by using the probe Kdn-BTP3, which forms a fluorescent precipitate upon hydrolysis. Using this method, they observed Kdnase activity both in the hyphae and extracellularly. They speculated that the fluorescent signal could originate from Kdnases in secretory vesicles bound to the plasma membrane or associated with the cell envelope. (12) In contrast to this work, our fluorescence readout comes from the covalent Kdnase-probe intermediate, which results in the direct visualization of the enzymes’ location. The *Af*Kdnase gene encodes a signal peptide and a secretion signal, (25) suggesting that the enzyme could be secreted or localized in the cell wall. While our findings cannot determine the exact location of the enzyme, they seem to indicate that Kdnases could be present at the hyphal surface, we cannot exclude intracellular staining but more work is required to prove this. (12) Nejatie et al.’s Kdn-BTP3 probe was internalized and interacted with intracellular Kdnase. Similarly, we hypothesize that our probe could also be internalized, achieving intracellular labeling either via the internalization of the probe clicked to alkyne-PEG_4_-AF_488_, or via internalization of both the probe and the click mixture containing alkyne-PEG_4_-AF_488_. The internalization of these molecules could occur through passive diffusion or via active transporters, as the *A. fumigatus* genome contains genes encoding several nucleotide sugar transporters, as suggested in a previous work. (25)

Finally, the role of *A. fumigatus* Kdnase in the fungal life cycle and in pathogenesis is not yet fully understood. However, it has been reported to play a critical role in fungal cell wall morphology and virulence. (15) To gain further insight, we monitored the germination of spores into hyphae in a media containing 0.3% yeast and malt extract, 0.5% peptone, and 1% glucose, supplemented with 200 µM **2e**,**3a-diF-9AzKdn**. We did not observe any differences in fungal morphology or growth rate under these conditions (supporting information S6-7). However, this experiment should be expanded by testing a broader range of concentrations and continuous addition of the inhibitor to more rigorously assess its potential effects.

## Conclusion

In this work, we developed difluoro-Kdn mechanism-based probes functionalized with azide and biotin tags for the selective inhibition, labeling and visualization of *Aspergillus fumigatus* Kdnases. These probes exhibited high selectivity and can efficiently label recombinant *Af*Kdnases at micromolar concentrations. Moreover, the azide-bearing probe followed by click chemistry enabled direct visualization of Kdnases in *Aspergillus fumigatus* mycelia, confirming its suitability for studies in complex biological matrices. Together, these results establish a versatile set of chemical tools for the direct detection and visualization of active retaining Kdnases in fungal systems. Beyond *A. fumigatus*, the probes offer opportunities to discover and characterize Kdnases in a broad range of fungal and bacterial species. Ultimately, this work provides a foundation for future studies aimed at elucidating the biological functions of Kdnases and assessing their potential as therapeutic targets.

## Materials and methods

### General

#### Synthesis and characterization of the Kdn ABPs

**The** synthesis and characterization data (^1^H-NMR, ^13^C-NMR, ^19^F-NMR, ESI-MS) can be found in supporting information S1.

#### Cloning, expression, and purification of *Aspergillus fumigatus* Kdnase (*Af*Kdnase) in *Escherichia coli*

The pSCodon1.2 expression vector was used to construct *Af*Kdnase fused to a C-terminal His-tag. The DNA fragment (*A. fumigatus* AFUA_4G13800) was elongated adding sequences overlapping the NdeI and XhoI sites of the linear pSCodon1.2 plasmid (see supporting information S5). The DNA fragment and the vector were assembled using the Gibson Assembly Master Mix according to manufacturer’s instructions. *E. coli* DH5α cells were transformed with pSCodon1.2_AFUA4G13800 plasmid. Cells were plated over LB-agar with ampicillin, colonies were grown at 37°C for 16 h. Plasmid was extracted from overnight LB broth-ampicillin cultures (37°C). *E. coli* BL21 cells were transformed with the extracted plasmid and plated over LB-agar plates supplemented with ampicillin. Pre-cultures (16 h, 37°C) were used to inoculate 50 mL of LB broth (1:50). Bacteria were grown at 37°C until OD_600_ 0.8. Heterologous protein expression was induced by the addition of isopropyl-β-D-thiogalactopyranoside to a final concentration of 1 mM. After 16 h of induction at RT, bacteria were collected by centrifugation (4,000 × g, 10 min) and stored at −80°C.

His-tagged full length *Af*Kdnase was purified under denaturing conditions. Bacterial pellets were suspended in a buffer 50 mM NaH_2_PO_4_, 300 mM NaCl, 10 mM imidazole, pH 8, supplemented with EDTA-free complete protease inhibitor cocktail (Roche Diagnostics) and 8 mg/mL lysozyme (Sigma) and incubated on ice for 30 min. The suspension was treated with 1 μL of benzonuclease (Pierce Universal Nuclease, 88700, ThermoFisher) and incubated at RT for 15 min. Bacteria were disrupted by sonication (10x 15 s pulses with 15 s holds on ice). The lysate was centrifuged (4.000 rpm, 30 min, 4°C), and the solvent of the soluble fraction was exchanged to a buffer 50 mM NaH_2_PO_4_, 300 mM NaCl, 10 mM imidazole, pH 8. The protein suspension was incubated with 1 mL of Ni^2+^-NTA agarose beads (Thermo Fisher Scientific) for 1 h at 4°C, the beads were washed x4 with 10 mL a buffer 50 mM NaH_2_PO_4_, 300 mM NaCl, 20 mM imidazole, pH 8. The protein was eluted with a buffer 50 mM NaH_2_PO_4_, 300 mM NaCl, 250 mM imidazole, pH 8. The fractions containing the protein were pooled together and concentrated using Spin-X UF 10 kDa filters (Corning). The protein concentration was determined by the BCA protein assay kit (Pierce™, Thermo 23225).

5’**GTTTAACTTTAAGAAGGAGATATA<u>CATATG</u>**atcaacgacccggccaagtcggccgccccctaccacgatgaattccctctcttccgcagcgccaa catggcctcgccagacaaactgtccaccggaatcggcttccactcgttccgcatccccgctgtggtccgcaccaccaccggacgcatcctcgccttcgccgagggccgccgacacaccaaccaagactttggcgacatcaaccttgtctacaagcgcaccaagacaaccgccaacaacggcgccagtccgtccgactgggagcctctccgggaagtcgtcggctccggcgccggcacctggggcaatccgaccccggtcgtcgacgacgacaacaccatctacctgtttctctcgtggaacggcgccacctacagccagaacggcaaggacgtcctccccgacggcacggtcaccaagaaaatcgactccacctgggaaggccgccgacacctgtacctcaccgagtcccgagacgacggcaacacctggtccaagcccgtcgatctcaccaaggaactgaccccggacggctgggcctgggacgcggtcggccccggcaacggcatccgcctcacgaccggcgagctggtcatccccgccatgggccgcaacatcatcgggcgcggcgcgccgggcaaccgcacgtggagcgtgcagcggctgtccggggccggggcggaggggacgatcgtgcagacgcccgacgggaagctgtaccgcaatgaccggcccagccagaaggggtaccggatggtggcgcgcgggacgctggagggcttcggggcgtttgccccggacgctgggctgccggacccggcgtgccagggatcggtgctgcggtataacagcgatgcgccggcgcggacgatctttctgaattccgcgtcggggacgagtcggcgggcgatgcgcgtgcggatcagctacgatgcggacgcgaagaagttcaactacgggcgcaagctggaggatgccaaggtcagcggggcgggtcatgaagggggttattcgagtatgaccaagacgggggattacaagattggggcgttggtggagagcgatttcttcaatgatggcactggcaagaattcgtatcg ggcgatcatctggaggagattcaatctgtcgtggatcctgaatggtcctaacaat**<u>CTCGAG</u>CACCACCACCACCACCACTGAGATC**-3’

Overlapping fragments highlighted in **red**.

#### Strains and growth conditions

*Aspergillus fumigatus* strains Af293.1, CEA10, ATCC46645 and A1258 were grown on potato dextrose agar plates at 37°C for 3 days. The spores were harvested and counted as described previously (26) and subsequently grown in 0.3%, w/v, yeast extract (Sigma Aldrich) and malt extract (Sigma Aldrich); 0.5%, w/v, peptone (Sigma Aldrich); and 20 mM glucose medium or minimal saline medium (MM; 6 g/L NaNO_3_, 1.5 g L-1 KH_2_PO_4_, 0.5 g/L, KCl, 0.5 g/L MgSO_4_·7H_2_O, 0.2 g/L Vishniac) (26) under shaking at 37°C for 3 days to obtain the mycelia. Mycelium was harvested via centrifugation before labeling in an Eppendorf tube at 150 rpm.

#### *Af*Kdnase activity assays

The activity of the recombinantly expressed *Af*Kdnase was determined by measuring cleavage of the synthetic fluorescent substrate (4-methylumbelliferyl)-3-deoxy-D-glycero-D-galacto-non-2-ulopyranosidonic acid (4MU-Kdn) (synthesis described in the supporting information 1). The reactions were set up in Corning® 96-well plates (Merck) by adding 160 µM 4MU-Kdn, recombinant enzyme (0.1 μM), and sodium acetate buffer pH 4 for a total volume of 100 µL. The plates were incubated at 37°C for 10 min followed by addition of 100 µL of stop solution (0.1% glycine, 25% ethanol, pH 10). To test the Kdnase activity of *A. fumigatus* washed mycelium was incubated with 4MU-Kdn (200 µM) for 2.5 h at RT. An aliquot of the supernatant (100 µL) was added to a Corning® 96-well plates (Merck) containing 100 µL of stop solution. The amount of 4-methylumbelliferone released from 4MU-Kdn was determined using a CLARIOstar Plus plate reader at excitation and emission wavelengths of 365 and 450 nm, respectively.

#### Kinetic assay

Michaelis-Menten parameters for the *Af*Kdnase were measured using 4MU-Kdn. Similar to previously reported. (9) Briefly, reaction mixtures (100 µM) were incubated at 37 °C for 10 min prior to the addition of the *Af*Kdnase. The progress of the reaction was continuously monitored for 30 min using a fluorescence spectrophotometer equipped with a temperature controller set to 37 °C and excitation and emission wavelengths of 365 and 450 nm, respectively. The kinetic parameters were determined from 5 initial rate measurements using a substrate concentration range of 100-600 μM, every measurement was done in duplicate. The rate versus substrate concentration data were fitted to the Michaelis-Menten equation using GraphPad.

#### Inhibition assays

Enzyme solutions were incubated with varying concentrations of the probes for 1 hour at RT. Kdnase activity was assessed by adding 4MU-Kdn (200 µM, 100 µL total volume on well), followed by a 30-minute incubation at RT. Reactions were stopped by the addition of 100 µL of the stop solution (0.1% glycine, 25% ethanol, pH 10). Fluorescence was measured using a Synergy H1 plate reader (Biotek) at excitation and emission wavelengths of 365 and 450 nm, respectively, with a gain setting of 1124. The apparent IC_50_ values were calculated using the Onesite-Fit logIC_50_ protocol in GraphPad.

To test inhibition of Kdnase in *A. fumigatus* mycelia, washed mycelia in sodium acetate buffer (50mM, pH 4), was incubated with 2e,3a-diF-9AzKdn (20 µM) for 2.5 h at RT. After this, 4MU-Kdn (200 µM) was added, and the reactions were incubated for 1 h at RT. An aliquot of the supernatant (100 µL) was added to a 96-well plate containing 100 µL of stop solution. The amount of 4-methylumbelliferone was measured with the plate reader as mentioned above.

#### Time-course study for *Af*Kdnase reactivation after incubation with probe 2e,3a-diF-9AzKdn

An enzyme solution (final concentration 10 nM) was incubated with **2e**,**3a-diF-9AzKdn** (final concentration 100 µM) in a 50 mM sodium acetate buffer pH 4 for 60 minutes at room temperature (RT). The solution was resuspended in 200 µL of Ni-NTA beads and incubated at room temperature (RT) for 30 minutes. The beads were washed three times with 400 µL of cold PBS. The Ni-NTA beads were then resuspended in 800 µL of the sodium acetate buffer described previously. At specific time points, 50 μL of the Ni-NTA suspension was mixed with 50 μL of 4MU-Kdn (100 μM final concentration) and incubated for 10 minutes. The supernatant was added to a well containing 100 μL of stop solution. Fluorescence was measured with a Fluostar reader (340 ex., 490 em.).

#### Labeling of recombinant *Af*Kdnase

Aliquots of the *Af*Kdnase enzyme (0.1 µM) were incubated with or without the probes in 50 mM sodium acetate buffer pH 4 for 1 hour at RT. As a control, samples containing heat-inactivated enzymes (boiled at 95 °C for 5 minutes) were included. Probe 2,3-diF-Neu9biotin was used to evaluate the selectivity of the enzyme towards Kdn-substrates. We also included bacterial neuraminidases from *Prevotella timonensis* (*Pt*NanH1 and *Pt*NanH2) to test the specificity of the probes. After incubation, reactions were stopped by adding 3×Laemmli buffer, followed by boiling at 95 °C for 5 minutes. The final reaction volume was 30 µL. A total of 15 µL from each sample was loaded onto a 10% Bis-Tris gel and electrophorized at 80–120 V using MOPS buffer (pH 7.0). For Western blotting, proteins were transferred to a PVDF membrane via electroblotting. The membrane was then blocked with 5% skimmed milk in PBS containing 0.5% Tween-20 (PBST). After blocking, the membrane was washed once with 1% skimmed milk in PBST for 5 minutes. It was then incubated with streptavidin-HRP (1:10000 in 1% skimmed milk) for 1 hour at RT. After washing with 1% skimmed milk in PBST and PBS (5 minutes each), the membrane was treated with ECL Western substrate for signal detection.

#### Fluorescent labeling and visualization of Kdnase in *A. fumigatus* mycelia by fluorescence microscopy

*A. fumigatus* ATCC46645 mycelium was incubated with 50 µM **2e**,**3a-diF-9AzKdn** for 2 hours at 37 °C, in sodium acetate buffer 50 mM, pH 4. Mycelium were washed with sodium acetate buffer, PBS and H_2_O three times each. Mycelium was resuspended in 100 µL of click mixture (1 µM alkyne-PEG_4_-AF_488_ (Jena Bioscience, Jena, Germany), 0.5 mM CuSO_4_, 2.5 mM sodium ascorbate in H_2_O) and incubated at RT for 2 h. Controls of mycelium without **2e**,**3a-diF-9AzKdn** were also treated with the click mixture (click controls), or just H_2_O (negative controls). After incubation, mycelium was washed three times with 200 µL of PBS, followed by fixation with 200 µL of 4% PFA in PBS for 30 min. Mycelium was washed three times with 200 µL of PBS and mounted on glass slides using 10 µL of ProLong Diamond Antifade Mountant (P36961, Thermo Fisher). Images of hyphae were collected on Invitrogen™ EVOS™ M5000 Imaging System, 40X and 100X, light exposure 0.05; and Leica® light transmitted microscope.

## Authors contributions

E.I.A.M.: Writing - Original Draft, Review and Editing, Conceptualization, Methodology, Resources, Investigation, Validation and Formal Analysis. J.N.: Culture of the fungi. H.C.: Conceptualization, Review and Editing, Formal Analysis, Supervision. T.W.: Conceptualization, Writing, Review and Editing, Formal Analysis, Supervision, Funding acquisition and Project administration.

## Acknowledgements

We thank Dr. Karin Strijbis (Utrecht University) for generously providing the recombinant *Pt*NanH1 and *Pt*NanH2 neuraminidases.

## Supporting Information

### 1. Supporting Information S1

Synthesis and characterization data (^1^H-NMR, ^13^C-NMR, ^19^F-NMR, ESI-MS).

#### Synthesis and characterization

All reagents were purchased from Sigma, Thermo Fisher Scientific and Biosynth and were used without further purification. Organic solvents were dried over pre-activated (2 h, 400 °C) 4 Å molecular sieves for 24 h prior to use. Glassware for anhydrous reactions was flame-dried and cooled under a nitrogen atmosphere. Thin layer chromatography (TLC) was performed on aluminium-backed Silica TLC Plates F254 (Silicycle, Canada) and detected by UV (254 nm or 365 nm) where applicable and by dipping in 10% sulfuric acid in ethanol, *p*-anisaldehyde sugar stain or ceric ammonium molybdate stain, followed by heating. Analytical thin layer chromatography (TLC) was performed on glass-backed TLC plates pre-coated with silica gel (60G, F_254_). Column chromatography was carried out using silica gel (40-63 μm; VWR chemicals) or C18-reversed phase silica gel (40-63 μm; Sigma). Solvents were evaporated at maximum 40 °C under reduced pressure. *N*-Acetylneuraminic Acid Aldolase from microorganisms (NANA aldolase) was purchased from Sigma (A6680). Dowex®, 1×8, 100-200 mesh resin was purchased from Thermo Fisher.

Nuclear magnetic resonance (NMR) spectra were recorded on a 400 MHz Agilent spectrometer (400 and 101 MHz) or a 600 MHz Bruker Avance Neo spectrometer (600 and 125 MHz). Chemical shifts are reported in parts per million (ppm) relative to residual solvent peak. Mass spectra were recorded by ESI on a Bruker micrOTOF-QII mass spectrometer.

#### 6-Azido-6-deoxy-D-mannose (2)

This compound was synthesized as previously reported in the literature. (1)

The ^1^H and ^13^C spectra of this compound were identical to previously reported spectra.

#### (4-methylumbelliferyl)-3-deoxy-D-glycero-D-galacto-non-2-ulopyranosidonic acid (4MU-Kdn)

This compound was synthesized as previously reported in the literature. (2,3)

The ^1^H and ^13^C spectra of this compound were identical to previously reported spectra.

#### 9-Azido-3-fluoro-D-glycero-D-galacto-non-2-ulopyranosidonic acid (3)

This compound was synthesized using a an adapted version of a previously published protocol for the synthesis of 3F-Neu9Ac. (4) Briefly, 6-azido-6-deoxy-**D**-mannose (422 mg, 2.3 mmol) was dissolved in 22 mL of PBS 0.05 M, pH 7.2, supplemented with 0.05% NaN_3_. βF-pyruvate (275 mg, 1.9 mmol) was added, followed by 20 mg of the NANA aldolase enzyme. The reaction was shaken at 37 °C, and monitored by ^19^F-NMR. Over a period of 5 days, 2 more equivalents of βF-pyruvate were added. The enzyme was precipitated by addition of 20 mL of cold ethanol, the mixture was centrifuged at 4 krpm for 5 min, and the supernatant was collected and evaporated. The residue was purified by ion exchange chromatography (Dowex®, 1×8, 100-200 mesh resin, formate form eluting with H_2_O to formic acid 2 M). Lyophilization gave the product consisting of a mixture 3F equatorial/axial (2:1), as a white solid with a 73% yield (490 mg, 1.7 mmol). ^1^H NMR (400 MHz, D_2_O) δ 4.75 (dd, *J* = 49.0, 2.3 Hz, 0.5H), 4.46 (dd, *J* = 49.5, 9.4 Hz, 1H), 3.95 – 3.81 (m, 3H), 3.80 – 3.67 (m, 3H), 3.60 – 3.53 (m, 1H), 3.48 (dd, *J* = 13.8, 3.0 Hz, 1H), 3.41 – 3.37 (m, 1H), 3.34 (dd, *J* = 13.1, 5.5 Hz, 1H). ^13^C NMR (101 MHz, D_2_O) δ 210.44, 187.93, 176.41, 166.13, 91.08, 89.94, 71.78, 70.87, 69.05, 66.96, 65.48, 58.49, 53.88, 42.15. ^19^F NMR (376 MHz, D_2_O) δ −199.97 (dd, *J* = 49.6, 13.6 Hz), −207.27 (dd, *J* = 49.7, 30.5 Hz), −228.18 – −230.45 (m).

#### 2,4,5,7,8-Penta-*O*-acetyl-9-azido-3-fluoro-D-glycero-D-galacto-non-2-ulopyranosonic acid methyl ester

To a solution of compound **3** (485 mg, 1.67 mmol) in CH_3_OH (25 mL), 1.0 mL of trifluoroacetic acid was added. The reaction was stirred at RT for 18 h. The solvent was evaporated and the residue was used for the next step without further purification. The crude methyl ester (510 mg, 1.57 mmol), a mixture of the equatorial/axial 3-fluoro isomers (2:1), was dissolved in 10 mL of dry pyridine at 0 °C under N_2_ atmosphere, then Ac_2_O (2.4 mL, 25.4 mmol) was added. The reaction mixture was left to reach room temperature (RT) and stir for 18 h. The mixture was diluted with 50 mL of ethyl acetate, and washed three times with 50 mL of a solution 0.1 M HCl, and two times with 50 mL of H_2_O. The organic layer was dried over Na_2_SO_4_, filtered and evaporated under reduced pressure. The residue was loaded onto a silica gel column and eluted with 10% ethyl acetate in PE → 30% ethyl acetate in PE to give the anomer **4** (314 mg, 0.59 mmol, 37%) and **5** (113 mg, 0.21 mmol, 14%) as white solids.

#### 2,4,5,7,8-Penta-*O*-acetyl-9-azido-3-fluoro(equatorial)-D-glycero-D-galacto-non-2-ulopyranosonic acid methyl ester (4)

R_f_ = 0.6 (EtOAc:PE, 1:1). ^1^H NMR (400 MHz, CDCl_3_) δ 5.50 (dt, *J* = 12.1, 9.4 Hz, 1H), 5.36 (dd, *J* = 4.2, 2.3 Hz, 1H), 4.99 (t, *J* = 9.9 Hz, 1H), 4.93 (ddd, *J* = 7.1, 4.2, 2.9 Hz, 1H), 4.56 (dd, *J* = 48.6, 9.3 Hz, 1H), 4.10 (dd, *J* = 10.3, 2.3 Hz, 1H), 3.83 (s, 3H), 3.80 – 3.73 (m, 1H), 3.34 (dd, *J* = 13.6, 7.1 Hz, 1H), 2.25 (s, 3H), 2.13 (s, 3H), 2.06 (s, 3H), 2.05 (s, 3H), 2.00 (s, 3H). ^13^C NMR (101 MHz, CDCl_3_) δ 170.12, 169.84, 169.46, 167.97, 77.30, 77.18, 76.98, 76.66, 72.43, 71.03, 70.65, 67.28, 66.06, 53.64, 49.89, 20.82, 20.77, 20.57, 20.54, 20.42. ^19^F NMR (376 MHz, CDCl_3_) δ −200.61 (dd, *J* = 48.6, 12.1 Hz). HRMS (ESI): *m/z* calcd for C_20_H_26_FN_3_O_13_+Na^+^ 558.1347; found: 558.1329.

#### 2,4,5,7,8-Penta-*O*-acetyl-9-azido-3-fluoro(axial)-D-glycero-D-galacto-non-2-ulopyranosonic acid methyl ester (5)

R_f_ = 0.55 (EtOAc:PE, 1:1). ^1^H NMR (400 MHz, CDCl_3_) δ 5.37 (dd, *J* = 3.7, 2.4 Hz, 1H), 5.32 – 5.29 (m, 1H), 4.97 (dd, *J* = 48.3, 2.0 Hz, 1H), 4.97 (ddd, *J* = 7.6, 3.7, 2.7 Hz, 1H), 4.13 – 4.05 (m, 1H), 3.88 (dd, *J* = 13.6, 2.7 Hz, 1H), 3.84 (s, 3H), 3.40 (dd, *J* = 13.6, 7.7 Hz, 1H), 2.19 (s, 3H), 2.14 (s, 3H), 2.08 (s, 3H), 2.07 (s, 3H), 2.00 (s, 3H). ^13^C NMR (101 MHz, CDCl_3_) δ 170.12, 77.30, 77.18, 76.98, 76.66, 72.90, 72.04, 67.56, 63.36, 53.60, 50.03, 20.83, 20.59, 20.54, 20.49. ^19^F NMR (376 MHz, CDCl_3_) δ −208.04 (dd, *J* = 48.4, 27.8 Hz). HRMS (ESI): *m/z* calcd for C_20_H_26_FN_3_O_13_+Na^+^ 558.1347; found: 558.1322.

#### 4,5,7,8-Tetra-*O*-acetyl-9-azido-3-fluoro(equatorial)-D-glycero-D-galacto-non-2-ulopyranosonic acid methyl ester (6)

A solution of hydrazine acetate (56 mg, 608 µmol) in 2 mL of dry CH_3_OH was added to a solution of compound **4** (63 mg, 126 µmol) in 2 mL of dry CH_2_Cl_2_ at 0 °C under argon atmosphere. The reaction was stirred at 0 °C for 7 h. The mixture was concentrated under reduced pressure, resuspended in 50 mL of ethyl acetate, and washed three times with 50 mL of a solution 0.5 M HCl, two times with 50 mL of a 5% solution of NaHCO_3_, 50 mL of H_2_O and 50 mL of brine. The organic layer was dried over Na_2_SO_4_, filtered and evaporated under reduced pressure. The residue was loaded onto a silica gel column and eluted with PE → 50% ethyl acetate in PE to give compound **6** (20 mg, 41 µmol, 32%) as a colorless syrup. R_f_ = 0.7 (EtOAc:PE, 1:1). ^1^H NMR (400 MHz, CDCl_3_) δ 5.49 (dt, *J* = 12.1, 9.5 Hz, 1H), 5.27 (d, *J* = 9.5 Hz, 2H), 5.20 (ddd, *J* = 7.9, 6.3, 3.0 Hz, 1H), 4.95 (ddd, *J* = 10.1, 9.4, 0.5 Hz, 1H), 4.85 (dd, *J* = 49.3, 9.5 Hz, 1H), 4.30 (dd, *J* = 10.4, 2.1 Hz, 1H), 3.92 (s, 3H), 3.42 (dd, *J* = 13.5, 3.0 Hz, 1H), 3.27 (dd, *J* = 13.5, 6.3 Hz, 1H), 2.12 (s, 3H), 2.10 (s, 3H), 2.03 (s, 3H), 2.01 (s, 3H). ^13^C NMR (101 MHz, CDCl_3_) δ 169.90, 169.81, 169.72, 169.70, 167.36, 167.34, 93.46, 93.25, 88.20, 86.24, 71.35, 71.16, 69.51, 68.60, 66.97, 66.67, 66.60, 54.32, 50.81, 29.21, 22.30, 20.76, 20.66, 20.62, 20.57, 20.49, 14.02. ^19^F NMR (376 MHz, CDCl_3_) δ −208.04 (dd, *J* = 48.6, 28.3 Hz). HRMS (ESI): *m/z* calcd for C_18_H_24_FN_3_O_12_+Na^+^ 516.1241; found: 516.1229.

#### 4,5,7,8-Tetra-*O*-acetyl-9-azido-3-fluoro(axial)-D-glycero-D-galacto-non-2-ulopyranosonic acid methyl ester (6)

A solution of hydrazine acetate (128 mg, 1.39 mmol) in 3 mL of dry CH_3_OH was added to a solution of compound **5** (63 mg, 126 µmol) in 2 mL of dry CH_2_Cl_2_ at 0 °C under argon atmosphere. The reaction was stirred at 0 °C for 6 h. The mixture was concentrated under reduced pressure, resuspended in 50 mL of ethyl acetate, and washed three times with 50 mL of a solution 0.5 M HCl, two times with 50 mL of a 5% solution of NaHCO_3_, 50 mL of H_2_O and 50 mL of brine. The organic layer was dried over Na_2_SO_4_, filtered and evaporated under reduced pressure. The residue was loaded onto a silica gel column and eluted with 20% ethyl acetate in PE → 25% ethyl acetate in PE to give compound **7** (109 mg, 221 µmol, 81%) as a transparent solid. R_f_ = 0.6 (EtOAc:PE, 1:1). ^1^H NMR (400 MHz, CDCl_3_) δ 5.41 – 5.30 (m, 1H), 5.29 (t, *J* = 2.5 Hz, 2H), 5.26 (dd, *J* = 10.2, 1.6 Hz, 1H), 4.92 (dd, *J* = 49.7, 2.1 Hz, 1H), 4.24 (dd, *J* = 10.1, 1.9 Hz, 1H), 3.88 (s, 3H), 3.61 (dd, *J* = 13.5, 2.5 Hz, 1H), 3.38 (dd, *J* = 13.4, 7.0 Hz, 1H), 2.13 (s, 3H), 2.12 (s, 3H), 2.06 (s, 3H), 2.02 (s, 3H). ^13^C NMR (101 MHz, CDCl_3_) δ 170.77, 170.14, 170.03, 169.43, 167.39, 94.21, 88.18, 77.30, 77.18, 76.98, 76.67, 71.27, 70.31, 70.15, 69.63, 67.50, 64.01, 53.72, 50.84, 20.98, 20.65, 20.62, 20.56, −0.04. ^19^F NMR (376 MHz, CDCl_3_) δ −200.54 (dd, *J* = 49.2, 12.1 Hz). HRMS (ESI): *m/z* calcd for C_18_H_24_FN_3_O_12_+H^+^ 494.1422; found: 494.1109.

#### 4,5,7,8-Tetra-*O*-acetyl-9-azido-2,3-difluoro(eq,eq)-D-glycero-D-galacto-non-2-ulopyranosonic acid methyl ester (8)

Compound **6** (20 mg, 41 µmol) was dissolved in 3 mL of dry CH_2_Cl_2_ under argon atmosphere and cooled to −30 °C. DAST (10 µL, 76 µmol) was added dropwise and the resulting reaction mixture was stirred at −30 °C for 1 h. The reaction was quenched by addition of 100 µL of CH_3_OH, the solvent was evaporated and the residue was resuspended in 10 mL of CH_2_Cl_2_. The organic layer was washed twice with 10 mL of a solution 0.4 M HCl, an ice cold 1% NaHCO_3_ (aq.) solution, and twice with 10 mL of ice cold water. The organic layer was dried over Na_2_SO_4_, filtered and evaporated under reduced pressure. The residue was purified via silica gel column chromatography (PE → 50% EtOAc in PE). Compound **8** (12 mg, 24 µmol) was obtained with a yield of 40%. R_f_ = 0.7 (EtOAc:PE, 1:1) as a yellow syrup. ^1^H NMR (400 MHz, CDCl_3_) δ 5.57 (ddd, *J* = 16.7, 8.8, 6.7 Hz, 1H), 5.30 (dt, *J* = 7.6, 1.4 Hz, 1H), 5.25 (ddd, *J* = 7.6, 5.8, 2.9 Hz, 1H), 5.09 (ddd, *J* = 10.3, 8.8, 1.3 Hz, 1H), 4.84 – 4.62 (m, 1H), 4.54 (dt, *J* = 10.7, 2.2 Hz, 1H), 3.58 (dd, *J* = 13.6, 2.9 Hz, 1H), 3.29 (dd, *J* = 13.6, 5.8 Hz, 1H), 2.11 (s, 3H), 2.10 (s, 3H), 2.06 (s, 3H), 2.03 (s, 3H). ^13^C NMR (101 MHz, CDCl_3_) δ 169.73, 169.61, 169.54, 169.40, 164.40, 164.08, 90.90, 90.56, 88.99, 88.65, 77.30, 76.99, 76.67, 72.06, 72.04, 71.68, 71.62, 71.44, 71.38, 69.64, 66.63, 65.78, 65.72, 53.61, 50.54, 20.67, 20.55, 20.51. ^19^F NMR (376 MHz, CDCl_3_) δ −115.03 (t, *J* = 12.4 Hz), −197.94 (ddd, *J* = 47.5, 16.7, 13.7 Hz). HRMS (ESI): *m/z* calcd for C_18_H_23_F_2_N_3_O_11_+Na^+^ 518.1198; found: 518.1378.

#### 4,5,7,8-Tetra-*O*-acetyl-9-azido-2,3-difluoro(eq,ax)-D-glycero-D-galacto-non-2-ulopyranosonic acid methyl ester (9)

Compound **7** (28 mg, 57 µmol) was dissolved in 3 mL of dry CH_2_Cl_2_ under argon atmosphere and cooled to −30 °C. DAST (20 µL, 151 µmol) was added dropwise and the resulting reaction mixture was stirred at −30 °C for 2 h. The reaction was quenched by addition of 100 µL of CH_3_OH, the solvent was evaporated and the residue was resuspended in 10 mL of CH_2_Cl_2_. The organic layer was washed twice an ice cold 1% NaHCO_3_ (aq.) solution, and twice with 10 mL of ice cold water. The organic layer was dried over Na_2_SO_4_, filtered and evaporated under reduced pressure. The residue was purified via silica gel column chromatography (20% EtOAc in PE → 30% EtOAc in PE). Compound **9** (27 mg, 55 µmol) was obtained with a yield of 96%. R_f_ = 0.7 (EtOAc:PE, 1:1) as a yellow syrup. ^1^H NMR (400 MHz, CDCl_3_) δ 5.34 (dt, *J* = 6.9, 2.1 Hz, 1H), 5.31 – 5.23 (m, 2H), 5.22 – 5.02 (m, 2H), 4.14 (dd, *J* = 9.9, 2.3 Hz, 1H), 3.91 (d, *J* = 0.5 Hz, 3H), 3.65 (dd, *J* = 13.6, 3.1 Hz, 1H), 3.30 (dd, *J* = 13.6, 6.0 Hz, 1H), 2.13 (s, 3H), 2.13 (s, 3H), 2.09 (s, 3H), 2.03 (s, 3H). ^13^C NMR (101 MHz, CDCl_3_) δ 169.73, 169.61, 169.54, 169.40, 164.40, 164.08, 90.90, 90.56, 88.99, 88.65, 77.30, 76.99, 76.67, 72.06, 72.04, 71.68, 71.62, 71.44, 71.38, 69.64, 66.63, 65.78, 65.72, 53.61, 50.54, 20.67, 20.55, 20.51. ^19^F NMR (376 MHz, CDCl_3_) δ −124.45 (dt, *J* = 11.2, 2.3 Hz), −215.84 (ddd, *J* = 50.4, 25.0, 10.9 Hz). HRMS (ESI): *m/z* calcd for C_18_H_23_F_2_N_3_O_11_+Na^+^ 518.1198; found: 518.1378.

#### 9-Azido-2,3-difluoro(eq,eq)-D-glycero-D-galacto-non-2-ulopyranosidonate (2e,3e-diF-9AzKdn)

Compound **8** (27 mg, 57 µmol) was dissolved in 2 mL of dry CH_3_OH, under argon atmosphere and Na_2_CO_3_ (6.8 mg, 111 µmol) was added. The reaction mixture was stirred at RT for 2 h. Next, the solvent was evaporated under reduced pressure. The residue was redissolved in an aqueous solution of NaOH 0.1 M (pH 11), and the mixture was stirred for 1 h at RT. The resulting solution was lyophilized, resuspended in H_2_O, and loaded onto a C18 silica gel reversed-phase column (gradient: 0% → 30% CH_3_CN in H_2_O). Compound **2eq**,**3eq-diF-Kdn** (27 mg, 56 µmol) was obtained in quantitative yield. R_f_ = 0.5 (EtOAc/CH_3_OH/H_2_O, 7:2:1). ^1^H NMR (400 MHz, D_2_O) δ 4.43 (ddd, *J* = 49.5, 13.7, 8.0 Hz, 1H), 4.31 – 4.13 (m, 1H), 4.10 (d, *J* = 10.6 Hz, 1H), 3.87 – 3.76 (m, 1H), 3.73 – 3.58 (m, 2H), 3.54 – 3.42 (m, 1H), 3.34 (dd, *J* = 13.3, 6.3 Hz, 1H). ^13^C NMR (151 MHz, D_2_O) δ 169.28, 111.66, 93.61, 74.43, 72.81, 68.95, 68.00, 67.64, 67.59, 53.89, 33.12. ^19^F NMR (376 MHz, D_2_O) δ −112.11 (t, *J* = 13.3 Hz), −200.63 (ddd, *J* = 49.5, 16.9, 13.2 Hz). HRMS (ESI): *m/z* calcd for C_9_H_13_F_2_N_3_O_7_+Na^+^ 336.0619; found: 336.0462.

#### 9-Azido-2,3-difluoro(eq,ax)-D-glycero-D-galacto-non-2-ulopyranosidonate (2e,3a-diF-9AzKdn)

Compound **9** (12 mg, 24 µmol) was dissolved in 2 mL of dry CH_3_OH, under argon atmosphere and Na_2_CO_3_ (6.8 mg, 111 µmol) was added. The reaction mixture was stirred at RT for 2 h. Next, the solvent was evaporated under reduced pressure. The residue was redissolved in an aqueous solution of NaOH 0.1 M (pH 11), and the mixture was stirred for 1 h at RT. The resulting solution was lyophilized, resuspended in H_2_O, and loaded onto a C18 silica gel reversed-phase column (gradient: 0% → 30% CH_3_CN in H_2_O). Compound **2eq**,**3ax-diF-Kdn** (6.0 mg, 19 µmol) was obtained with a yield of 79%. R_f_ = 0.6. (EtOAc/CH_3_OH/H_2_O, 7:2:1). ^1^H NMR (400 MHz, D_2_O) δ 5.04 (dt, *J* = 51.0, 2.8 Hz, 1H), 3.99 – 3.88 (m, 1H), 3.88 – 3.84 (m, 1H), 3.79 (td, *J* = 9.8, 1.5 Hz, 1H), 3.76 – 3.70 (m, 1H), 3.66 – 3.60 (m, 1H), 3.53 (dd, *J* = 13.2, 2.7 Hz, 1H), 3.36 (dd, *J* = 13.2, 6.2 Hz, 1H). ^13^C NMR (151 MHz, D_2_O) δ 169.07, 105.64, 90.00, 89.88, 88.66, 73.67, 73.65, 71.14, 71.02, 69.34, 68.00, 65.28, 65.26, 53.67, 42.83, 18.51. ^19^F NMR (376 MHz, D_2_O) δ −121.81 (d, *J* = 11.6 Hz), −217.11 (ddd, *J* = 51.0, 28.4, 11.4 Hz). HRMS (ESI): *m/z* calcd for C_9_H_13_F_2_N_3_O_7_+Na^+^ 336.0619; found: 336.0600.

#### 9-Biotin-2,3-difluoro(eq,eq)-D-glycero-D-galacto-non-2-ulopyranosidonate (2e,3e-diF-9bKdn)

**2eq**,**3eq-diF-9AzKdn** (2.5 mg, 8.0 µmol) was dissolved in 1000 µL degassed H_2_O. Next, a solution of alkyne-PEG_4_-biotin (4.4 mg, 9.6 µmol) was added to the mixture, followed by the addition of CuSO_4_ with 3 equivalents of THPTA ligand (final concentration 0.3 mM), and a freshly prepared solution of Na*-L*-ascorbate (final concentration of 0.7 mM). The reaction was stirred at RT in the dark for 16 h. The crude reaction mixture was purified by C18 reversed-phase silica column (H_2_O→50% CH_3_CN in H_2_O) and the product was obtained as a white solid with 68% yield (4.2 mg, 5.5 µmol). ^1^H NMR (600 MHz, D_2_O) δ 4.62 (d, *J* = 4.8 Hz, 1H), 4.57 – 4.23 (m, 2H), 4.23 – 4.00 (m, 2H), 3.85 – 3.58 (m, 17H), 3.54 (t, *J* = 5.2 Hz, 2H), 3.30 (t, *J* = 5.2 Hz, 2H), 3.25 (dt, *J* = 9.7, 5.2 Hz, 1H), 2.91 (dd, *J* = 13.0, 5.0 Hz, 1H), 2.73 – 2.53 (m, 1H), 2.18 (t, *J* = 7.3 Hz, 2H), 1.69 – 1.41 (m, 2H), 1.32 (s, 2H). ^13^C NMR (151 MHz, D_2_O) δ 233.01, 182.64, 100.19, 88.14, 74.21, 69.57, 69.40, 68.85, 62.43, 62.05, 55.37, 41.65, 39.72, 35.43, 27.84, 27.65, 25.11. ^19^F NMR (376 MHz, D_2_O) δ −110.30, −200.31 (d, *J* = 50.6 Hz). HRMS (ESI): *m/z* calcd for C_30_H_48_F_2_N_6_O_13_S+H^+^ 771.3046; found: 771.3017.

#### 9-Azido-2,3-difluoro(eq,ax)-D-glycero-D-galacto-non-2-ulopyranosidonate (2e,3a-diF-9bKdn)

**2eq**,**3ax-diF-9AzKdn** (2.5 mg, 8.0 µmol) was dissolved in 1000 µL degassed H_2_O. Next, a solution of alkyne-PEG_4_-biotin (4.4 mg, 9.6 µmol) was added to the mixture, followed by the addition of CuSO_4_ with 3 equivalents of THPTA ligand (final concentration 0.3 mM), and a freshly prepared solution of Na*-L*-ascorbate (final concentration of 0.7 mM). The reaction was stirred at RT in the dark for 16 h. The crude reaction mixture was purified by C18 reversed-phase silica column (H_2_O→50% CH_3_CN in H_2_O) and the product was obtained as a white solid with 78% yield (4.8 mg, 6.2 µmol). ^1^H NMR (600 MHz, D_2_O) δ 4.57 – 4.27 (m, 2H), 4.17 (d, *J* = 2.3 Hz, 1H), 3.68 (dt, *J* = 5.9, 2.2 Hz, 1H), 3.67 – 3.59 (m, 10H), 3.55 (q, *J* = 4.6 Hz, 2H), 3.31 (dt, *J* = 9.1, 5.0 Hz, 2H), 3.28 – 3.22 (m, 1H), 2.92 (dd, *J* = 12.8, 4.9 Hz, 1H), 2.83 (td, *J* = 2.4, 1.0 Hz, 1H), 2.71 (dd, *J* = 13.1, 4.1 Hz, 1H), 2.25 – 2.08 (m, 2H), 1.71 – 1.43 (m, 2H), 1.39 – 1.25 (m, 2H). ^13^C NMR (151 MHz, D_2_O) δ 176.94, 165.37, 79.28, 75.96, 73.48, 69.64, 69.59, 69.56, 69.43, 69.41, 68.87, 68.68, 68.62, 65.28, 62.07, 60.23, 57.91, 55.45, 55.37, 39.80, 39.71, 38.90, 35.45, 35.43, 27.85, 27.66, 27.64, 25.11. ^19^F NMR (376 MHz, D_2_O) δ −120.83, −216.52. HRMS (ESI): *m/z* calcd for C_30_H_48_F_2_N_6_O_13_S+H^+^ 771.3046; found: 771.3245.

### 2. Supporting Information S2

Activity of *Af*Kdnase. Determination of K_m_.

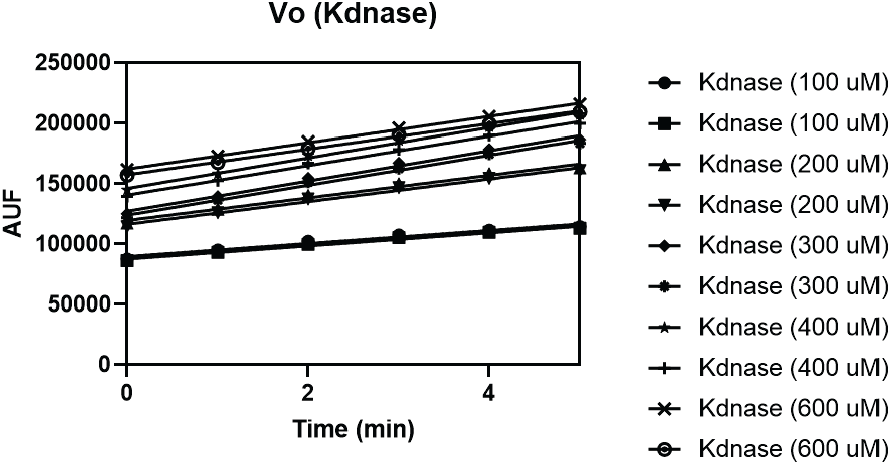

### 3. Supporting Information S3

Full-size gel.

### 4. Supporting Information S4

Fluorometric assay monitoring Kdnase activity in mycelia of different *A. fumigatus* strains.

### 5. Supporting Information S5

Plasmid map of *Aspergillus fumigatus* Kdnase (*Af*Kdnase) transformed into *Escherichia coli* BL21 (DE3).

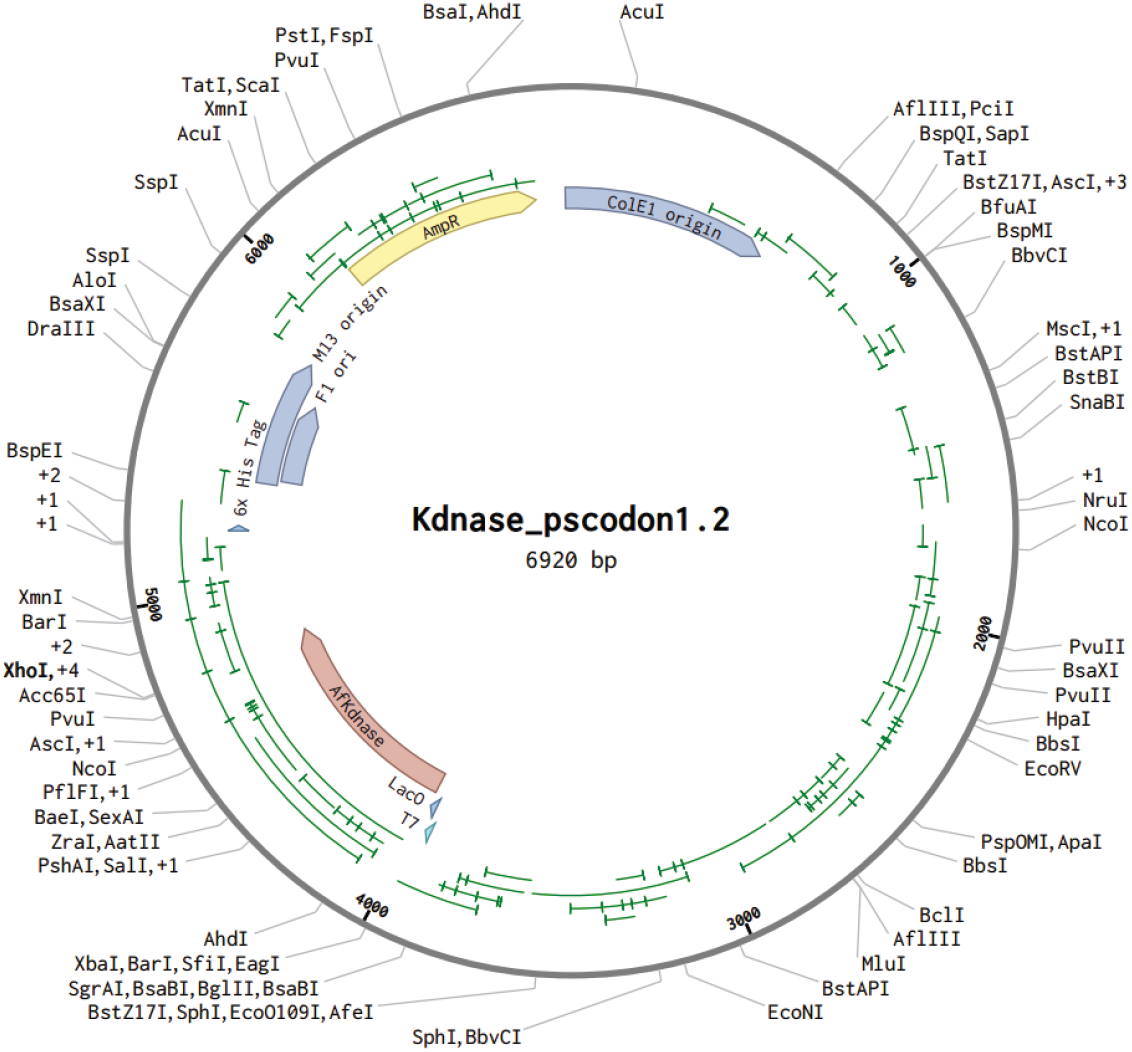

### 6. Supporting Information S6

Growth curve of *A. fumigatus* ATCC46645 in the presence of probe diF-Kdn9Az.

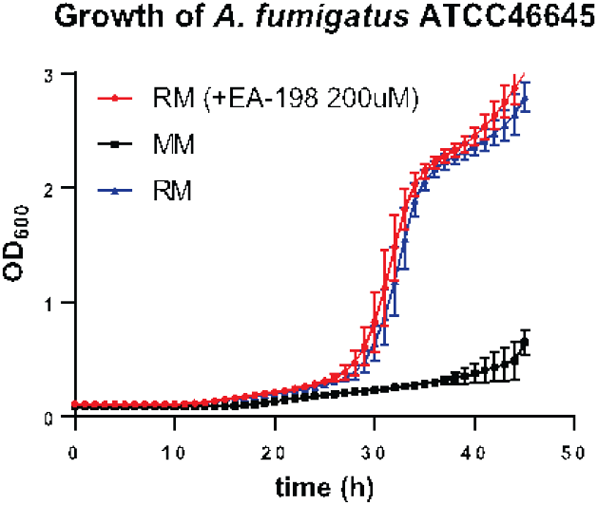

### 7. Supporting Information S7

Microscopy images. Phenotype of *Aspergillus fumigatus* ATCC46645 mycelia after 45 h incubation at 37 °C with and without **2**,**3-diF-9AzKdn** probe.

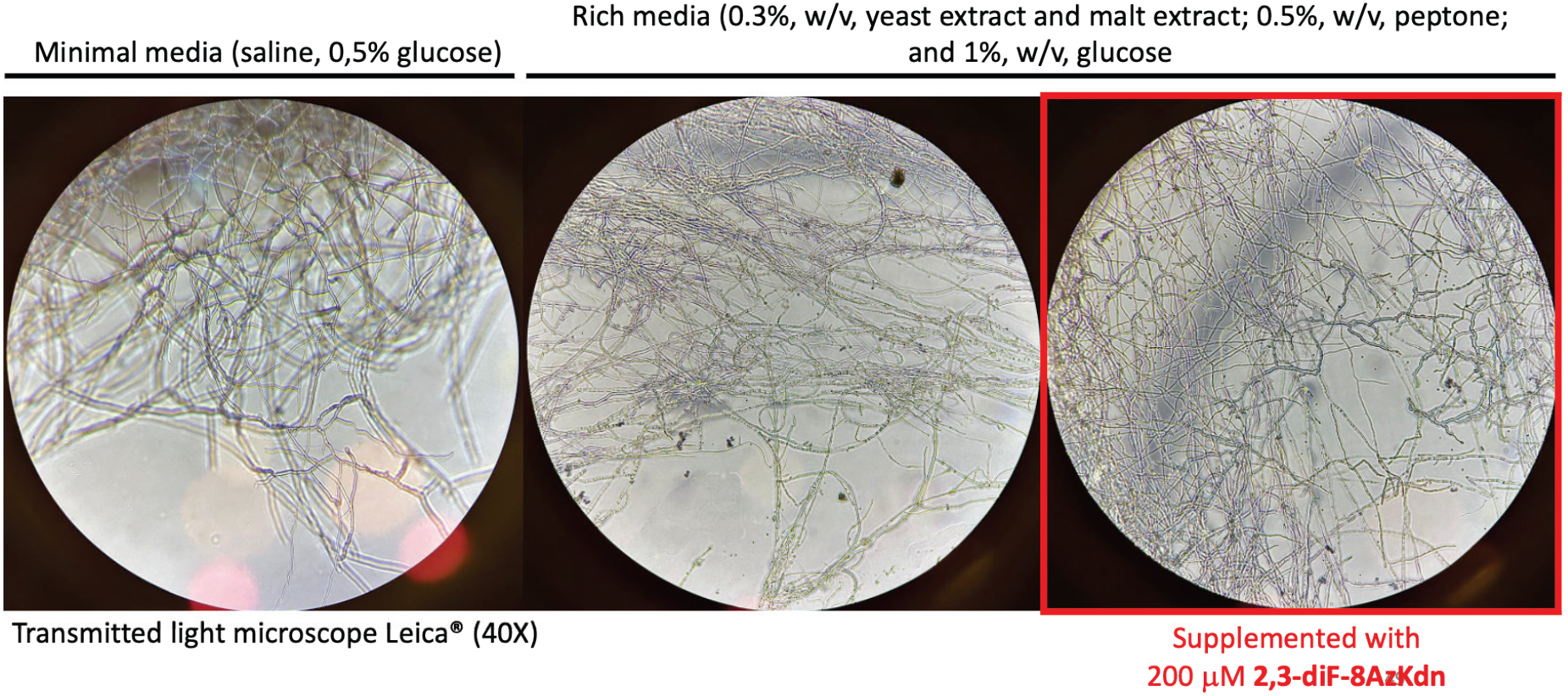

